# Strain-dependent disease progression and necrotizing granuloma formation induced by virulent *Mycobacterium avium* complex strains in a murine model

**DOI:** 10.1101/2025.08.03.668372

**Authors:** Haruka Hikichi, Shiho Omori, Hajime Nakamura, Shintaro Seto, Koji Furuuchi, Kozo Morimoto, Minako Hijikata, Naoto Keicho

## Abstract

*Mycobacterium avium* complex (MAC) is the leading cause of non-tuberculous mycobacterial pulmonary disease (NTM-PD), a chronic infection with a heterogeneous clinical course. Although murine models of MAC-PD have been developed, reproducing the key features of progressive human disease, including diverse pathological features and therapeutic sensitivities, remains challenging. In this study, we evaluated five clinical MAC strains, including a newly identified highly virulent isolate, NBRC112750, in immunocompetent BALB/c mice. Among these, FKJ-1 and NBRC112750 induced progressive pulmonary infection with increasing bacterial burdens and extensive lung involvement by 25 weeks post-infection. Notably, both strains led to the formation of necrotizing granulomas resembling those observed in *M. tuberculosis*-infected C3HeB/FeJ mice. These lesions featured neutrophilic infiltration, foamy macrophages, and collagen encapsulation. Using FKJ-1 strain, we also established an inhalation infection model, in which low-dose exposure reproduced necrotizing granulomas. Despite *in vitro* drug susceptibility, FKJ-1 infection exhibited poor response to standard therapy, highlighting strain-dependent variability in treatment efficacy. These findings establish a murine model that reflects both key pathological and therapeutic aspects of MAC-PD and provides a valuable platform for investigating MAC pathogenesis and evaluating novel therapies.

## Introduction

Pulmonary infection caused by non-tuberculous mycobacteria (NTM), referred to as NTM pulmonary disease (NTM-PD), is an increasingly prevalent chronic respiratory condition worldwide (1–3). Among NTM species, *Mycobacterium avium* complex (MAC), complising *M. avium* and *M. intracellulare*, is the leading causative agent of NTM-PD in several countries, including Japan (2, 4–6).

NTM-PD, including MAC-PD, presents a heterogeneous clinical spectrum. Two primary radiological phenotypes are recognized: the fibrocavitary (FC) type and the nodular bronchiectatic (NB) type, each associated with distinct demographics and disease courses (4, 7). The FC type, characterized by cavitary lesions predominantly in the upper lobes, is more prevalent among men with underlying PDs, including prior pulmonary tuberculosis and chronic obstructive PD. In contrast, the NB type typically affects postmenopausal women without a smoking history. The FC type is associated with a poorer treatment response and worse prognosis, whereas the NB type has a higher risk of NTM-PD recurrence following treatment (8). Despite these differences, both phenotypes share a common pathological feature: granulomatous inflammation throughout the airways (9, 10). Thus, granuloma formation is implicated in human MAC-PD pathogenesis. This peribronchial granulomatous inflammation is believed to contribute to airway wall damage and bronchial distortion, eventually leading to irreversible airway structural changes such as bronchiectasis.

The introduction of macrolide-based combination therapy, typically inducing clarithromycin (or azithromycin), ethambutol, and rifampicin, has significantly improved treatment success compared to earlier regimens (4, 7, 8). However, recurrence and persistent infection remain common, and treatment outcomes vary among patients, suggesting that host–pathogen interactions play a critical role in disease progression and therapeutic response (11, 12).

To better understand the pathogenesis of MAC-PD and to develop more effective treatments, several murine models of pulmonary MAC infection have been established (12, 13). However, existing models, particularly those using immunocompetent mice, rarely replicate key features of progressive human disease, such as the formation of necrotizing granulomas. Notably, C3HeB/FeJ mice infected with certain MAC strains could develop necrotizing granulomas (14), similar to those observed in the same mice following *M. tuberculosis* infection (15). However, another study failed to replicate these findings under same conditions (16).

We previously isolated clinical MAC strains from patients with MAC-PD and demonstrated that pulmonary infection in immunocompetent mice results in increased or sustained bacterial burdens and granulomatous lesion development (17). To expand our collection of clinical MAC strains relevant to the pathological features of MAC-PD, we screened additional isolates obtained from public microbiological repositories. Among them, one strain fulfilled our selection criteria by exhibiting increasing bacterial burdens and inducing pulmonary granuloma formation in mice. Given the chronically progressive nature of MAC-PD, we hypothesized that long-term infection in a murine model could recapitulate the disease progression and the pathological heterogeneity observed in patients. In this study, we evaluated five clinical MAC strains, including a newly identified, highly virulent isolate, using long-term infection models in BALB/c mice. Our aim was to determine whether strain-specific virulence contributes to differences in disease progression, lesion development, and variability in treatment response. Through this approach, we identified MAC strains capable of inducing chronic infection and necrotizing granulomas. Moreover, we observed strain-dependent differences in therapeutic responses, supporting the establishment of a murine model that more accurately reflects the pathological spectrum of MAC-PD.

## Results

### Virulence of the *M. intracellulare* strain NBRC112750 in mice

We collected clinical MAC strains from three public microbiological repositories in Japan: the Japan Collection of Microorganisms, the Gene bank project of the National Agriculture and Food Research Organization, and the Biological Resource Center of the National Institute of Technology and Evaluation. Strains were selected and evaluated in a mouse infection model based on the following criteria: (I) isolation from patients with MAC-PD, including specimens such as sputum or bronchoalveolar lavage fluid; (II) the ability to induce pulmonary granulomatous lesions following intranasal infection in mice; and (III) sustained or increasing mycobacterial burdens in the lungs at 8 weeks postinfection (p.i.). We screened MAC strains as described previously (17), and confirmed the virulence of the selected strains. Among the 18 MAC strains tested, only NBRC112750 met all these criteria. At 8 weeks p.i., mice infected with NBRC112750 exhibited peribronchial granulomas with increased the mycobacterial burdens in the lungs (Supplementary Figure 1).

### Disease progression in pulmonary MAC-infected mouse models

We previously isolated four clinical MAC strains that, when administered intranasally to BALB/c mice, led to the development of granulomatous lesions and increased or sustained mycobacterial burdens in the lungs (17). In this study, we evaluated disease progression in mouse models infected with five MAC strains, namely FKJ-1, FKJ-2, FKJ-5, FKJ-8, and NBRC112750, over a 24-week infection period. Among them, FKJ-1, FKJ-2, and NBRC112750 are *M. intracellulare*, while FKJ-5 and FKJ-8 are *M. avium*. MAC strains were intranasally administered to BALB/c mice at a dose of 10^6^ CFU. All strains demonstrated a progressive increase in lung mycobacterial burdens by 8 weeks p.i., which continued throughout the infection. Paticularly, FKJ-1 and NBRC112750 exceeded 10^8^ CFU in the lungs by 25 weeks p.i (Figure 1). Histological analysis revealed that all strains induced the formation of peribronchial granulomatous lesions at 8 weeks p.i. (Figure 2), as previously described (17). By 20 weeks p.i., lungs infected with FKJ-1 and NBRC112750 exhibited extensive inflammatory consolidation across lung lobes and the emergence of necrotizing granulomas with central necrotic cores, which eventually developed into well-defined necrotizing granulomas. Among other strains, excluding FKJ-1 and NBRC112750, although FKJ-2 and FKJ-5 demonstrated increased bacterial burdens and granuloma formation, disease progression over time was limited. In contrast, FKJ-8 exhibited an initial increase in bacterial burden; however, the overall bacterial load remained lower than that of FKJ-1 and NBRC112750 throughout the infection. Furthermore, the pathological changes induced by FKJ-8 did not progress to necrotizing granuloma formation, even during long-term infection. Owing to these differences in both bacterial dynamics and pathological progression, subsequent experiments focued on comparative analyses of the FKJ-1 and FKJ-8 strains.

**Figure 1.**
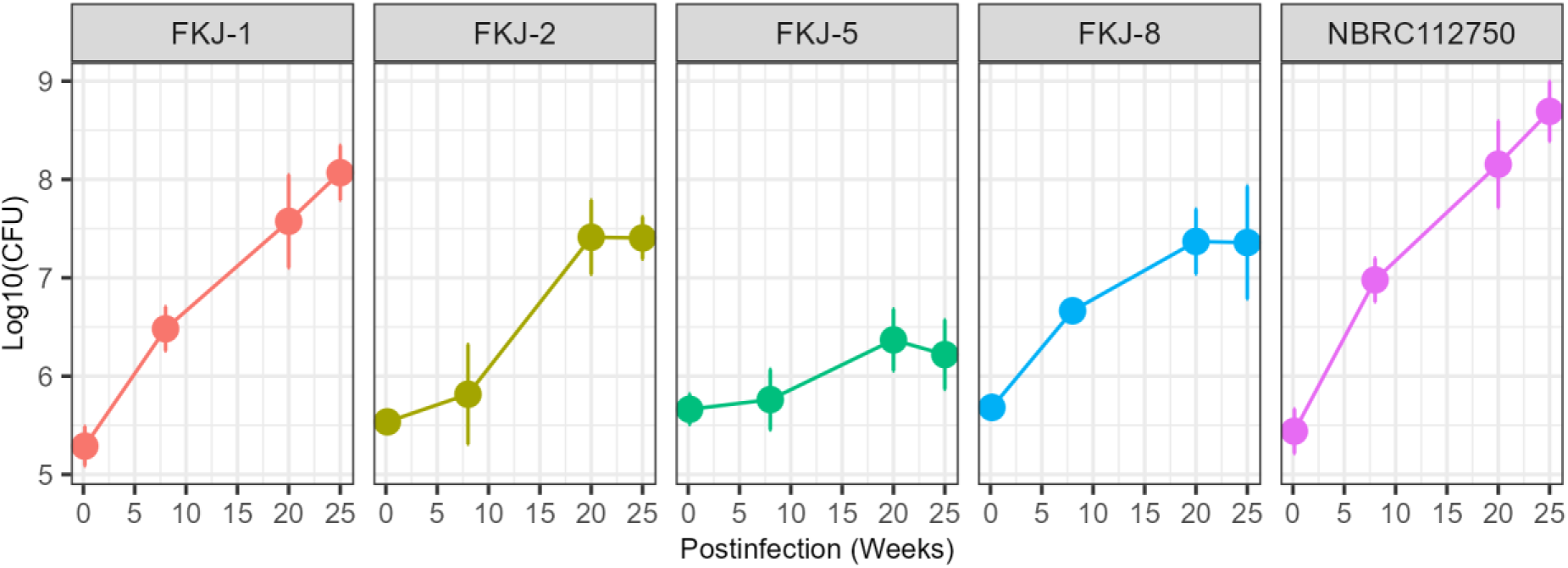
Mycobacterial burdens in infected lungs over a 25-week infection period. BALB/c mice were intranasally infected with 10^6^ CFU of the indicated *M. avium* complex (MAC) strains and monitored for 25 weeks. At each time point, the mean with standard deviation (SD) of CFU counts from six mice are shown. Multiple comparisons were conducted using Tukey–Kramer post hoc test. Statistical comparisons are presented in Supplementary Table 2.

**Figure 2.**
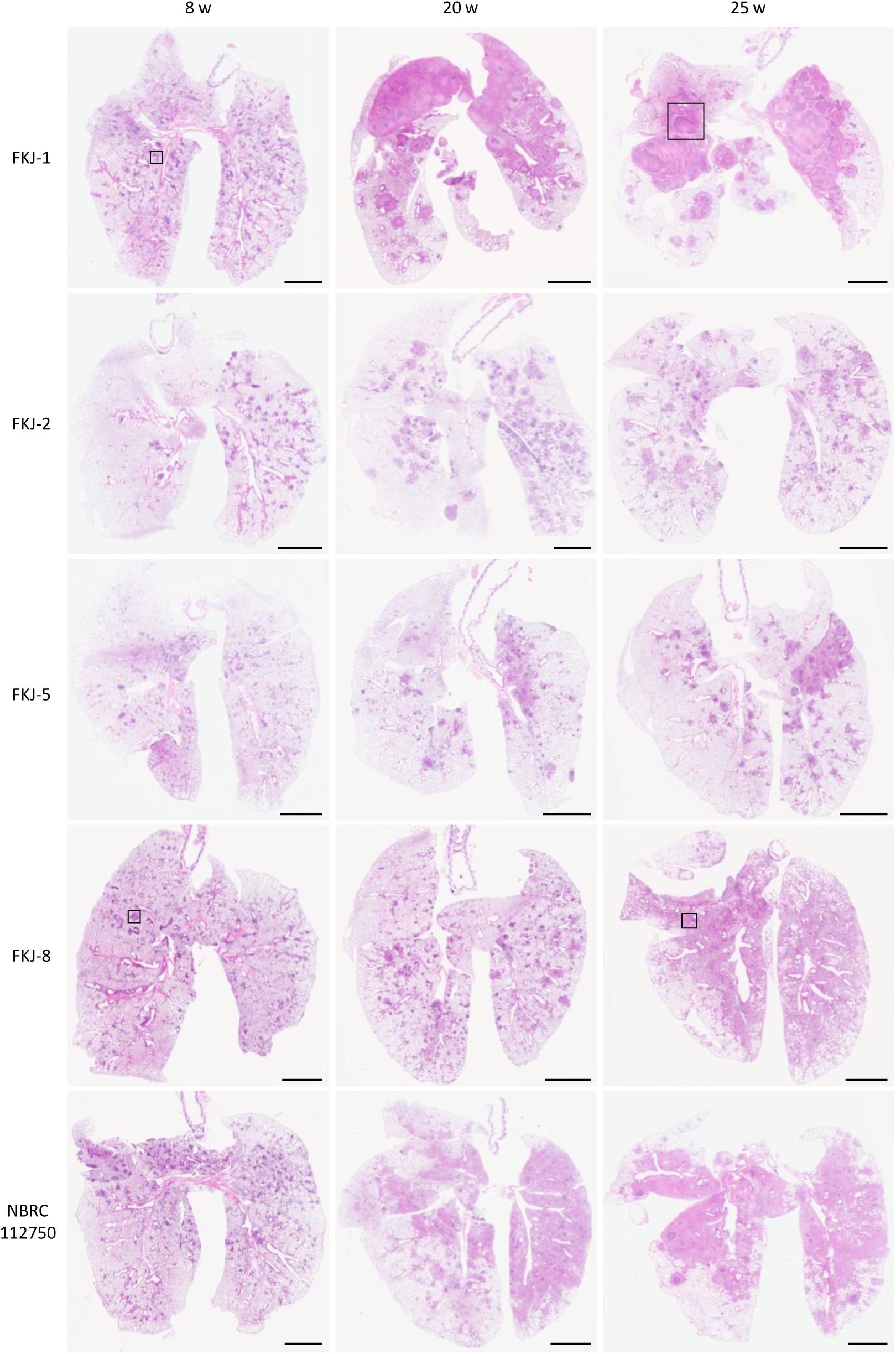
Development of mycobacterial granulomatous lesions in mouse lungs. Representative hematoxylin and eosin (H&E)-stained images of whole lung lobes from mice infected with the indicated MAC strains at respective time points. Boxed areas are shown at higher magnification in Figure 3. Scale bar, 2.5 mm.

### Necrotizing granulomas in FKJ-1-infected mouse lungs

We characterized granulomas in the lungs of MAC-infected BALB/c mice (Figure 3). Although both FKJ-1 and FKJ-8 strains induced peribronchial granulomas at 8 weeks p.i., FKJ-1-infected lungs developed necrotizing granulomas by 25 weeks p.i (Figure 3A-D). These lesions closely resembled those observed in *M. tuberculosis*-infected C3HeB/FeJ mice, as previously reported (18–22). Ziehl-Neelsen staining revealed the presence of replicating mycobacteria within epithelioid macrophages surrounding the necrotic cores. Masson’s trichrome staining showed that necrotizing granulomas in the lungs of FKJ-1-infected mice at 25 weeks p.i. were well-organized and encapsulated by collagen fibers. In contrast, other granulomas exhibited collagen deposition but lacked evident encapsulation (Figure 3E–H).

**Figure 3.**
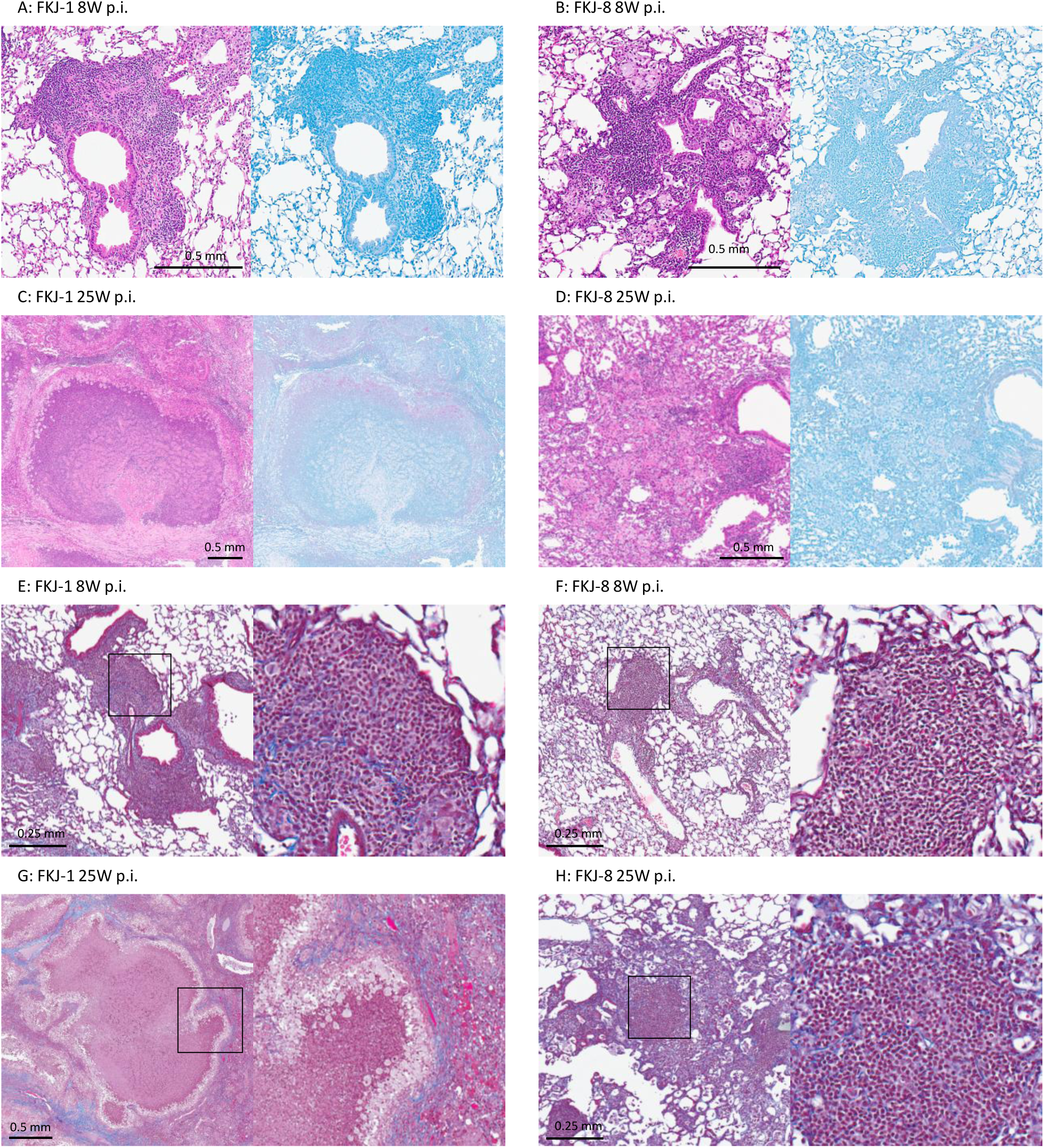
Enlarged images of granulomatous lesions in MAC-infected mouse lungs. H&E- and Ziehl–Neelsen (ZN)-stained sections from granulomatous lesions in mouse lungs infected with the FKJ-1 strain at 8 weeks postinfection (p.i) (A), FKJ-8 strain at 8 weeks p.i. (B), FKJ-1 strain at 25 weeks p.i. (C), and FKJ-8 strain at 25 weeks p.i (D). (E-H) Masson’s trichrome-stained sections showing collagen distribution in granulomatous lesions from mice infected with FKJ-1 or FKJ-8 strains at 8 or 25 weeks p.i. Right panels show the corresponding enlarged views of the boxed areas in left panels.

To further investigate the cellular composition of FKJ-1-induced necrotizing granulomas, we examined neutrophil accumulation within the lesions. Neutrophil infiltration is observed in necrotizing granulomas of patients with tuberculosis and MAC-PD, as well as in *M. tuberculosis*-infected C3HeB/FeJ mice (9, 18, 20–26). Immunostaining using an anti-Ly6G antibody, a neutrophil marker, revealed strong signals within the necrotic cores, indicating substantial neutrophil infiltration into the FKJ-1-induced necrotizing granulomas (Figure 4A). Additionally, immunostaining for foamy macrophage markers Plin2, Msr1, and Arg1, typically localized to the cells surrounding necrotic granulomas in *M. tuberculosis*-infected C3HeB/FeJ (18, 27), revealed their presence in FKJ-1-infected BALB/c mouse lungs (Figure 4B-D). These findings suggest that intranasal infection with the FKJ-1 strain over 24 weeks leads to necrotic granuloma development with a cellular composition similar to that observed in *M. tuberculosis*-infected C3HeB/FeJ mice.

**Figure 4.**
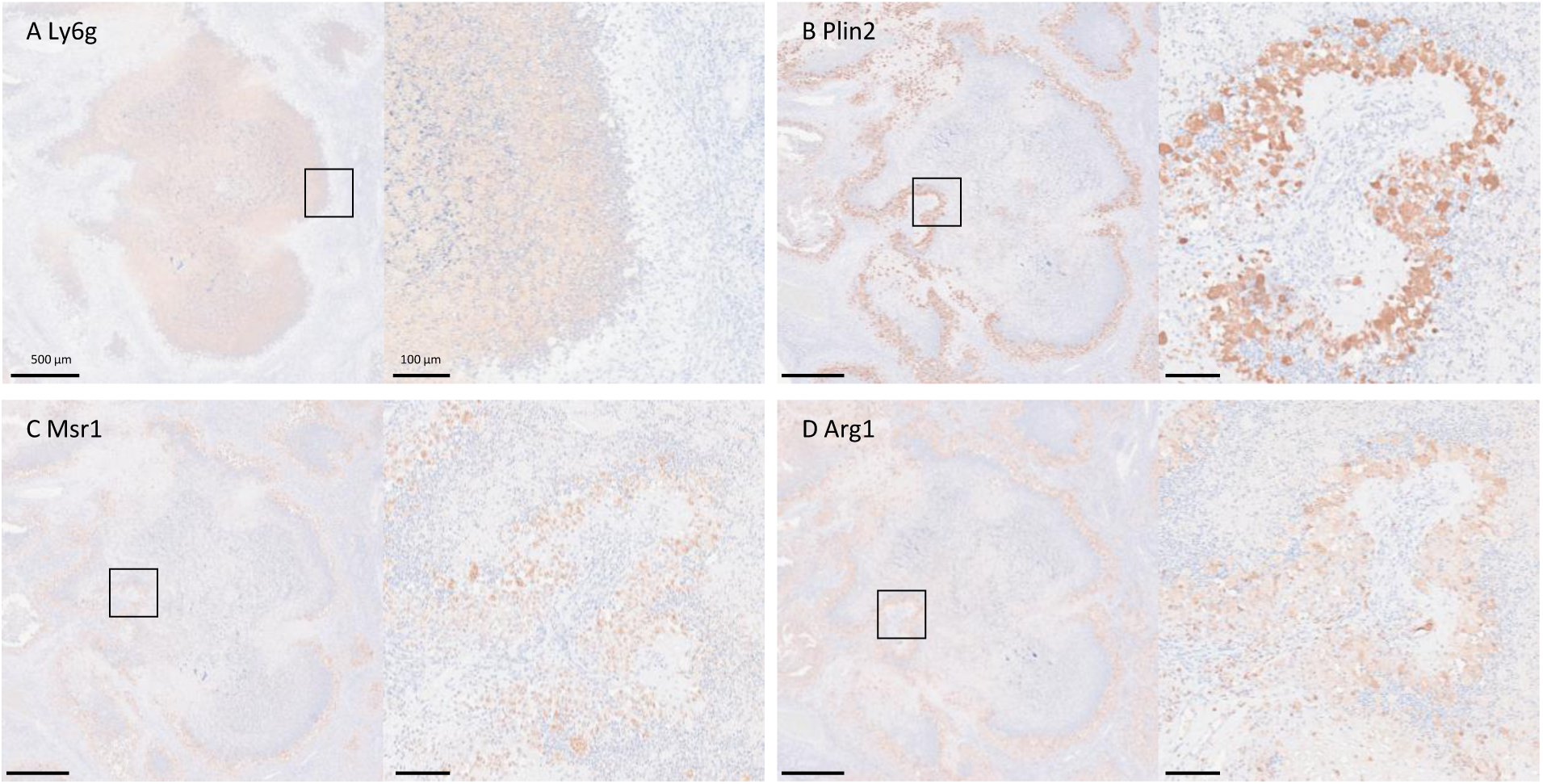
Localization of neutrophils and foamy macrophage markers in necrotizing granulomatous lesions in the lungs of mice infected with a MAC strain. Immunohistochemistry (IHC) analysis of necrotizing granulomatous lesions from the lungs of mice infected with the FKJ-1 strain at 25 weeks p.i., showing the neutrophil marker Ly6g (A) and foamy macrophage markers Plin2 (B), Arg1 (C), and Msr1 (D). Right panels display the corresponding enlarged images of the boxed areas in left panels.

### Establishment of a novel mouse model via aerosol exposure with a MAC strain

We assessed the virulence of the FKJ-1 strain in BALB/c mice via the aerosol route using an inhalation exposure system (Figure 5). Mice were infected with initial doses of 10^2^ (low), 10^3^ (moderate), and 10^4^ (high) CFU at one day p.i., and mycobacterial burdens and pathological features were monitored up to 24 weeks. Lung mycobacterial burdens exhibited clear dose-dependent dynamics during the infection (Figure 5A). Mice infected with low or moderate doses exhibited increased bacterial burdens for up to 8 or 16 weeks p.i., respectively, followed by a plateau. In contrast, high-dose infection resulted in sustained increases in lung mycobacterial burdens throughout the infection, indicating the development of a progressively chronic infection.

**Figure 5.**
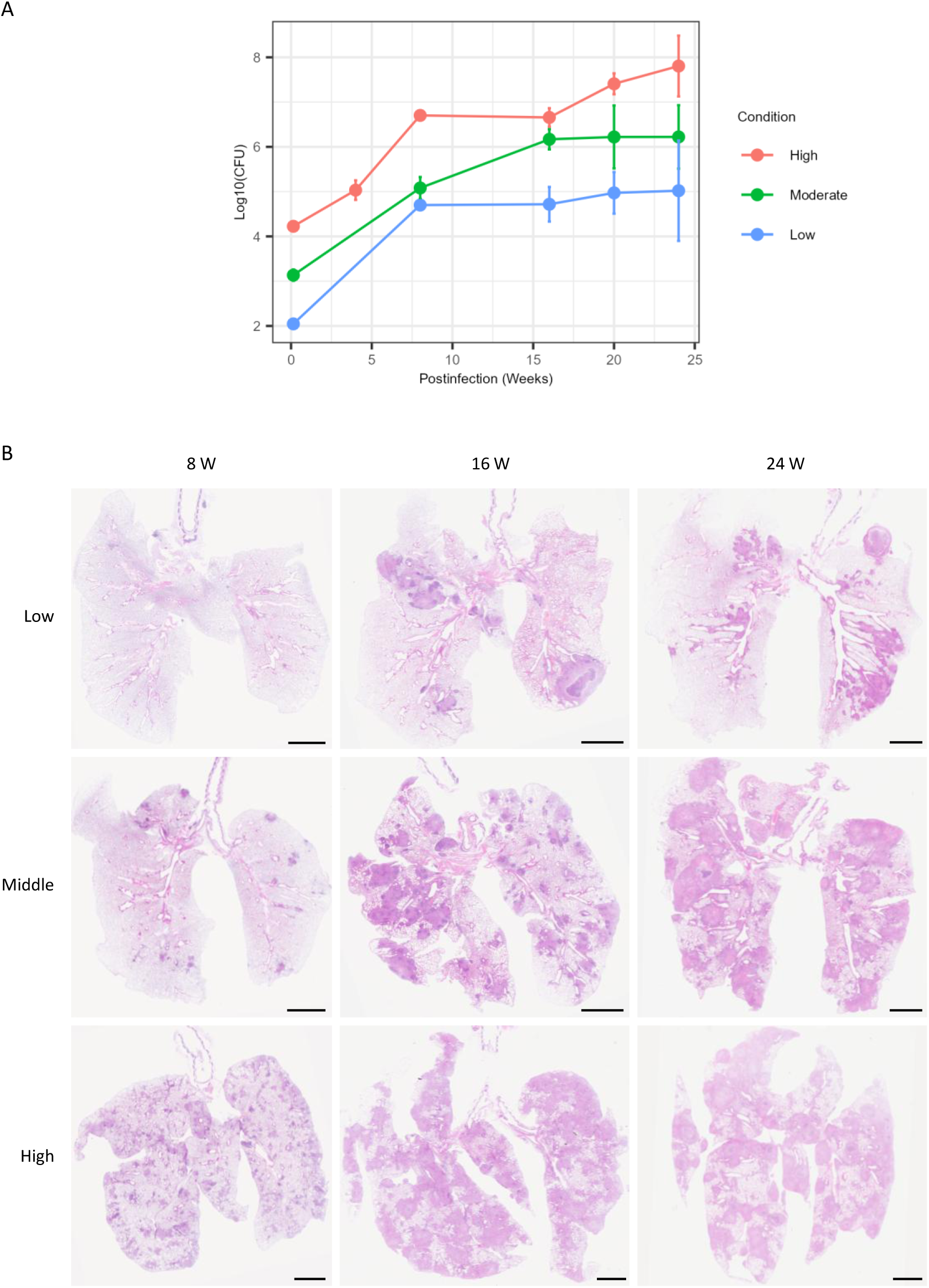
Aerosol infection of BALB/c mice with a MAC strain. Mice were infected with the FKJ-1 strain at three different doses via aerosol exposure and monitored for 24 weeks. (A) Mycobacterial burdens in mouse lungs. At each time point, the mean with SD of CFU counts from six mice are shown. Mice were infected at approximately 10^2^ CFU (Low), 10^3^ CFU (Moderate), and 10^4^ CFU (High) at one day p.i. Multiple comparisons were performed using Tukey– Kramer post hoc test. Statistical comparisons are presented in Supplementary Table 2. (B) Development of mycobacterial granulomatous lesions in mouse lungs. Each panel shows a representative H&E-stained image of whole lung lobes infected with the FKJ-1 strain at the indicated dose and time point.

Histopathologically, high-dose-infected mice exhibited numerous lesions across all lung lobes at 8 weeks p.i. In contrast, only a few lesions were found in the lungs of low-dose-infeced mice (Figure 5B). By 16 or 24 weeks p.i., moderate- or high-dose-infected micedeveloped granulomatous lesions throughout the lung lobes. At 24 weeks p.i., necrotic cores appeared within these granulomatous lesions. Notably, distinct necrotizing granulomas were also observed in low-dose-infected mice by 16 weeks p.i.

### Therapeutic efficacy in aerosol MAC-infected mouse models

We evaluated the therapeutic efficacy of the standard MAC-PD regimen, comprising clarithromycin, ethambutol, and rifampicin, in mouse models infected with MAC strains, FKJ-1 and FKJ-8 via aerosol exposure (Figure 6). The minimum inhibitory concentrations (MICs) of tested antibiotics against these strains are listed in Table 1. Mice were infected with a high dose (10^4^ CFU) of FKJ-1 or FKJ-8 at one day p.i. At 4 weeks p.i., chemotherapeutic treatment was continued for another 4 weeks. In FKJ-8-infected mice, mycobacterial burdens were reduced in drug-treated animals compared to controls at both 4 and 8 weeks p.i. In contrast, FKJ-1-infected mice exhibited only a modest reduction in bacterial burdens following treatment. At 8 weeks p.i., while the bacterial burdens continued to increase in control mice, those in drug-treated mice remained relatively stable (Adjusted *P* = 0.201). These findings suggest that FKJ-1 is less susceptible to the standard regimen *in vivo*, despite similar *in vitro* antibiotic sensitivity compared to FKJ-8.

**Figure 6.**
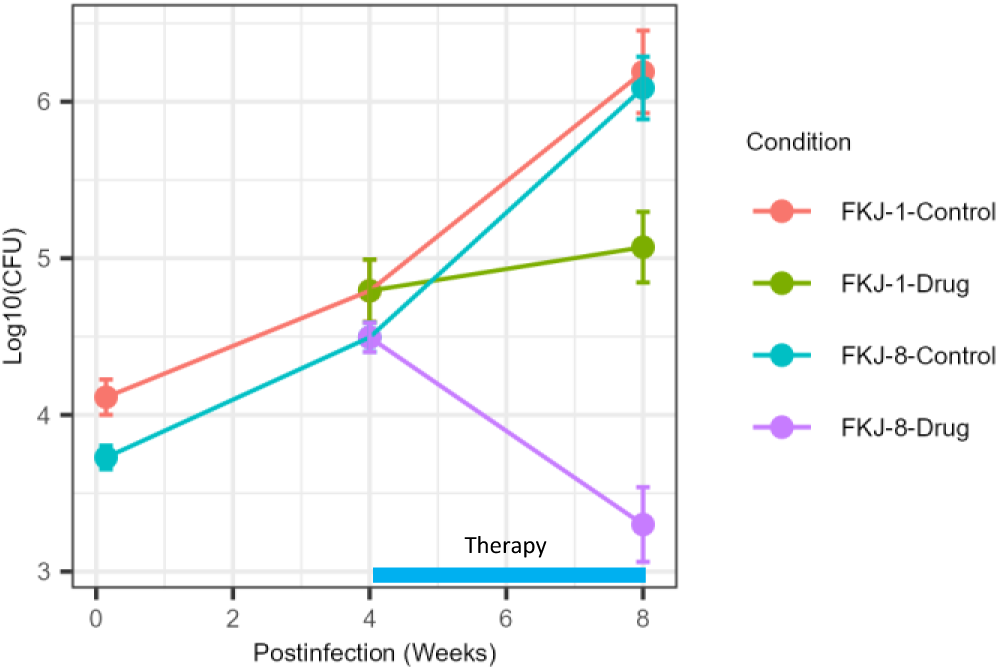
Therapeutic treatment of MAC-infected mice. Mice were infected with FKJ-1 or FKJ-8 strains via aerosol exposure. At 4 weeks p.i., infected mice were treated with a combination of rifampicin (10 mg/kg/mouse), ethambutol (100 mg/kg/mouse), and clarithromycin (100 mg/kg/mouse), administrated 5 days per week for 4 weeks. At each time point, the mean with SD of CFU counts from six mice are shown. Tukey–Kramer post hoc test was used for multiple comparisons. Statistical comparisons are presented in Supplementary Table 2.

**Table 1.** Minimum inhibitory concentrations of FKJ-1 and FKJ-8 strains.

| Strains | MIC (µg/mL) |  |  |
| --- | --- | --- | --- |
|  | Clarithromycin | Ethambutol | Rifampicin |
| FKJ-1 | 0.5 | 2 | 0.25 |
| FKJ-8 | 2 | 8 | 0.25 |

## Discussion

In this study, we identified a highly virulent MAC strain, NBRC112750, capable of inducing chronic and progressive lung infection in BALB/c mice. To evaluate the pathological heterogeneity of MAC-PD, we infected immunocompetent mice with five MAC strains, including NBRC112750 and four previously isolated strains (17), and monitored them over a prolonged period. Given that MAC-PD exhibits a variable clinical course, ranging from stable disease to progressive respiratory failure, we aimed to determine whether the pathological heterogeneity could be recapitulated in a murine model.

A remarkable outcome of chronic infection with FKJ-1 or NBRC112750 was the development of necrotizing granulomas in the lungs of BALB/c mice (Figure 2). By 20–25 weeks p.i., mice infected with either strain exhibited extensive inflammatory consolidation and necrotizing granulomas with central necrotic cores. In contrast, FKJ-2 and FKJ-8 induced non-necrotic granulomas, despite lung mycobacterial burdens exceeding 10⁷ CFU. The necrotizing granulomas formed in FKJ-1 or NBRC112750-infected BALB/c mice resembled those observed in C3HeB/FeJ mouse model of *M. tuberculosis* infection (18–22). We further characterized the features of necrotizing granulomas induced by FKJ-1 infection and observed that: (I) neutrophils accumulated within the necrotic cores; (II) foamy macrophage markers, including Plin2, Msr1, and Arg1, were expressed in epithelioid macrophages surrounding the necrotic areas; and (III) the granulomas were encapsuled by collagen fibers (Figures 3 and 4). These histopathological features were similar to those reported in *M. tuberculosis*-infected C3HeB/FeJ mice (18, 27). In that model, necrotizing granuloma formation has been attributed to reduced expression of *Sp110* and *Sp140*, which are negative regulators of type I interferon (IFN) expression (28–30). However, the development of similar necrotizing granulomas in BALB/c mice infected with MAC strains suggests a distinct underlying mechanism. Our findings imply that the magnitude or persistence of the host inflammatory response, rather than host genetics alone, may drive necrotizing pathology. The presence of abundant neutrophils and foamy macrophages harboring replicating bacilli supports the hypothesis that highly virulent MAC strains elicit a sustained, localized immune response that exceeds the threshold for tissue integrity, resulting in necrotizing granuloma development.

To further investigate host susceptibility, we infected C3HeB/FeJ mice with the FKJ-1 strain (Supplementary Figure 2). Notably, lower mycobacterial burdens were observed in the lungs of C3HeB/FeJ mice than in BALB/c mice, with no necrotizing granuloma development. These results suggest that C3HeB/FeJ mice, despite their susceptibility to *M. tuberculosis*, could control FKJ-1 infection. One possible explanation involves *Nramp1* (*Slc11a1*), an iron transporter in phagosomes that modulates the survival and replication of infected *M. avium* in macrophages (31). The functional allele of *Nramp1* is expressed in C3H mice, the parental strain of C3HeB/FeJ. In contrast, BALB/c and C57BL/6 mice harbor a nonfunctional *Nramp1* allele, which is associated with increased susceptibility to mycobacterial infection (32). These observations suggest that *Nramp1*-mediated restriction of intracellular mycobacteria may reduce the inflammatory response, thereby limiting lesion progression and preventing necrotizing granuloma formation. Overall, our results emphasize the complex interplay between bacterial virulence, host genetics, and the local immune microenvironment in determining granulomatous pathology.

We established a novel inhalation model of MAC infection using the FKJ-1 strain via an aerosol exposure system (Figure 5). Notably, low-dose infection with FKJ-1 induced necrotizing granuloma formation at 16 weeks p.i., occurring earlier than in moderate- or high-dose infections. This finding suggests that lesion development may depend more on the dynamics of host– pathogen interactions than on the initial bacterial load. In the low-dose infection model, gradual bacterial replication may allow for prolonged immune activation, resulting in sustained inflammation and the development of necrotizing granulomas. These observations suggest that chronic infection with virulent MAC strains also can drive necrotizing granulomatous inflammation.

Moreover, the differential therapeutic responses between FKJ-1 and FKJ-8 highlight the importance of identifying strain-specific virulence determinants and treatment susceptibility. Several studies have demonstrated the chemotherapeutic efficacy in MAC-infected mouse models (33–36). However, despite the similar *in vitro* antibiotic sensitivities, FKJ-1 infection resulted in persistent mycobacterial burdens *in vivo* during treatment (Figure 6). These findings suggest that our mouse model recapitulates the variable treatment responses observed in patients with MAC-PD, where certain clinical strains respond poorly to standard therapy despite confirming in vitro susceptibility (8, 37).

In this study, antimicrobial treatment was initiated at the early phase of infection, prior to necrotizing granuloma formation. Despite this early intervention, therapeutic outcomes differed between two strains, suggesting that early host–pathogen interactions may influence both *in vivo* drug efficacy and the trajectory of lesion development. Thus, this model offers a valuable platform for evaluating strain-dependent therapeutic efficacy and for developing more effective, tailored therapeutic strategies for MAC infection.

Overall, our findings highlight the complex interplay between bacterial virulence, host genetics, and persistent inflammatory responses in driving both necrotizing granuloma development and treatment variability in MAC-PD. The murine model presented here, which recapitulates key pathological and therapeutic aspects of the disease, represents a valuable platform for exploring host–pathogen interactions and optimizing more effective, strain-specific treatment strategies.

## Materials and Methods

### Ethics Statement

The animal experiment conducted in this study was approved by the Animal Care and Use Committee of The Research Institute of Tuberculosis (RIT) (Permit number: No. 2023-02). Animals were treated in accordance with the ethical guidelines of the RIT. The endpoints were set to determine whether the mice were imminently dying of MAC infection and/or required compassionate euthanasia: bodyweight loss > 20% of the initial bodyweight at the time of infection.

### Mice

BALB/c and C3HeB/FeJ mice were purchased from Japan SLC and Jackson Laboratory, respectively. C3HeB/FeJ mice were maintained in a filtered-air laminar-flow cabinet, and provided sterile bedding, water, and mouse chow at the RIT animal facility. Specific pathogen-free status was verified by testing sentinel mice housed within the colony. BALB/c mice aged 6 weeks and C3HeB/FeJ mice aged 6–10 weeks were transferred to the biosafety level (BSL) II or BSL III animal facility of the RIT.

### Infection of MAC strains

The clinical MAC strains, FKJ-1, FKJ-2, FKJ-5, FKJ-8 (17), and NBRC112750, isolated from human tracheal lavage fluid, were cultured in 7H9 medium supplemented with 10% Middlebrook ADC (BD Bioscience), 0.5% casamino acid and 0.05% Tween 80 (mycobacterial medium) (30) at 37°C. The NBRC112750 strain was provided by the National Institute of Technology and Evaluation. Single-cell suspensions were prepared as previously described (17) and stored at −80°C until use. Mice were infected with MAC strains either intranasally at a dose of 10^6^ CFU in 30 μL of saline or via the aerosol route using an infection exposure system (Glas-Col) at doses of 10^2^, 10^3^, or 10^4^ CFU to the lungs.

### Mycobacterial burdens in lungs and histological analysis

At each time point, infected mice were euthanized by exsanguination under anesthesia with 0.75 mg/kg medetomidine, 4.0 mg/kg midazolam, and 5.0 mg/kg butorphanol via the intraperitoneal route. To determine pulmonary mycobacterial burdens, infected lungs were homogenized using a FastPrep-24 5G instrument (MP Biomedicals). The resulting lung homogenates were serially diluted and plated on 7H10 or 7H11 agar plates supplemented with 10% Middlebrook OADC (BD Bioscience) and 0.5% glycerol. For histological analysis, the whole lung lobes from infected mice were fixed with 10% formalin in PBS for over 24 h at room temperature. Tissue sections were stained with hematoxylin and eosin (H&E), Ziehl–Neelsen (ZN) stain, or Masson’s trichrome for collagen etection. Immunohistochemistry (IHC) analysis was performed as previously described (9, 18, 27). Antibodies used in this study are listed in Supplementary Table 1. Stained sections were visualized using a NanoZoomer S60 slide scanner (Hamamatsu Photonics).

### Minimal inhibitory concentrations and chemotherapy

MICs were determined by inoculating MAC strains at 2.5 ×10^5^ CFU/mL in mycobacterial medium containing antibiotics in 96-well plates and incubating them for 5 days at 37°C. Clarithromycin, ethambutol, and rifampicin were purchased from Tokyo Chemical Industry. For chemotherapy, six mice infected with FKJ-1 or FKJ-8 at 4 weeks p.i. were orally administered a combination of clarithromycin (100 mg/kg), ethambutol (100 mg/kg), and rifampicin (10 mg/kg) in 100 μL of 0.5% methylcellulose solution (Fujifilm) 5 times per week for 4 weeks.

## Acknowledgments

This study was supported by the Emerging/Re-emerging Infectious Diseases Project of the Japan Agency for Medical Research and Development (JP23wm0225028, JP23gm1610013, JP23fk0108673, JP23fk0108674, JP23fk0108703, JP25fk0108730), and Grants-in-Aid for Scientific Research, Japan Society for the Promotion of Science (24K10229, 25K10370).

We thank Mr. Masanari Matsuda, Mr. Akihiro Hata, Ms. Miyako Seto, Ms. Mariko Ogasawara, and Ms. Akiko Ozawa of Department of Pathophysiology and Host Defense, RIT, for their technical supports. We also thank Dr. Shigeaki Hida and Dr. Isamu Ogawa of Nagoya City University, and Dr. Takemasa Takii of RIT, for their valuable consultation regarding chemotherapy in mice.

HH, SO, HN, performed experiments, analyzed data, and wrote and revised the manuscript. SS, MH, NK designed the project and analyzed data, and wrote and revised the manuscript. KF, KM provided the materials and revised the manuscript. All authors approved the manuscript.

The authors declare that the research was conducted in the absence of any commercial or financial relationships that could be construed as a potential conflict of interest.

